# Hydrodynamic cues enhance algal lipid production while sustaining biomass in motile species

**DOI:** 10.1101/2025.01.02.631047

**Authors:** Narges Kakavand, Anupam Sengupta

## Abstract

Achieving enhanced lipid yield without compromising biomass is one of the long-standing challenges in our quest to produce algal biofuel sustainably. Multiple factors, including temperature, nutrients and light conditions impact lipid production, however such lipid-enhancing strategies often lead to reduced biomass, thereby offsetting the total volume of lipid recovered. Hydrodynamic cues remain poorly studied, specifically in the context of lipid production in motile algae, concurrently with biomass generation and photo-physiology, a key fitness parameter. By imposing hydrodynamic cues to biophysically stress distinct strains of raphidophyte *Heterosigma akashiwo* at specific time points along the growth stages (indicating different nutritional states), we quantify the lipid production, alongside algal biomass and photo-physiology. Early induction (hydrodynamic cues implemented during the lag phase) and delayed induction (hydrodynamic cues implemented during the exponential phase) were studied. Delayed induction of hydrodynamic cues suppressed growth and photo-physiology without significant enhancement of lipid production, however, early induction allowed to significantly increase lipid content, up to 300%, without observable changes in biomass and photo-physiology. Based on this, we propose a hydrodynamic strategy for enhanced lipid production with sustained biomass and physiological fitness. This work presents hydrodynamic perturbation and its onset timing as tunable parameters to advance lipid production technologies across diverse motile species.

## I. INTRODUCTION

Biofuels, renewable and environmentally friendly alternatives to fossil fuels [1], have received renewed attention during the past decade in light of the exacerbated global warming [2–4]. Although awareness and demand for green bio-based alternatives to fossil fuels is at a peak, their energy efficiency and economic viability remain a moot point, preventing universal acceptance of biofuels [5]. Relatively lower energy yields [6], particularly in terrestrial plants which were the primary feedstock for biofuel production, has been a key hurdle, though microalgae has shown promise as the next generation of biofuel feedstock, thanks to their superior energy density, efficient growth, and ultimately higher yield as a sustainable source of energy [5, 7–9]. Beyond the biochemical advantages, microalgae offer operational benefits such as simplified cultivation, economic viability, and reduced environmental footprint in generation, further consolidating their preferred status over traditional plant-based feedstocks [8].

Microalgae, equipped with chlorophyll, play a crucial role in converting atmospheric carbondioxide to oxygen through photosynthesis. While microalgae hold exceptional promise as a feedstock for biofuel, sustainable production still faces obstacles due to the disproportionate energy requirements for biofuel production and extraction from algal feedstock [7, 10–12]. At the scale of a single cell, biofuels are produced in the form of cytoplasmic lipid droplets [5, 13], often under constraints of nutrients, light or mechanical perturbations which act as biophysical stressors.

Physiological stress due to altering light intensities [14–16], salinity [17–19], nutrient availability (nitrogen and phosphorus deficiencies) [13, 20, 21], temperature [22, 23], and mechanical compression [24] have been employed to drive alagal lipid production. Stressors, while promoting lipid accumulation, inadvertently compromise photophysiology and fitness (cell growth and population), suppressing overall biomass [25, 26]. In agreement with fundamental studies on the interactions between microalgae and their dynamic environments [13, 27, 28], the stressful conditions–a requirement algal biofuel production–lead to lower biomass, thus offsetting higher yields with reduced biomass. Two-stage cultivation systems, employing an initial stage for microalgal growth followed by induction of biofuel-regulating conditions, offer a way around [26], however they come at a higher production cost. Consequently, it becomes central to algal biofuel production that the balance between promoting growth and enhancing lipid production is attained at a cost-effective manner [29], and that the factors underpinning this delicate balance are well understood.

Among various microalgal species used in biofuel production, majority of them are non-motile due to their relatively higher lipid contents and predictable growth patterns, making them suitable for bioreactor cultivation [5, 30, 31]. More recently, there has been a growing recognition of motile species as potential biofuel sources, including *Chlamydomonas reinhardtii* [31], *Dunaliella tertiolecta* [32], *Dunaliella primolecta* [33], *Tetraselmis suecica* [34], *Isochrysis galbana* [35], and *Haematococcus pluvialis* [36]. Disruption of motility due to environmental factors may trigger physiological stress which could be leveraged for lipid production [13, 27, 28, 37]. Among motile species, raphidophytes offer distinct attributes which are amenable to biofuel production and extraction [38]. For instance, their rapid proliferation rates [28, 39], and absence of conventional cell wall resulting in direct access to lipid extraction (compared to algae with cell walls or thecas [40]) offer competitive advantages, ensuring energy-efficient extraction process [41]. Yet, raphidophytes remain laregly unexplored as a biofuel feedstock, due to paucity of research on their cultivation, lipid accumulation kinetics, and appropriate lipid extraction protocols, leaving open a major gap in our ability to fully harness these species for algal fuel generation [42–44].

*Heterosigma akashiwo*, a well known raphidophyte, produces neutral lipids under nutrient limitations, with strain-specific variation in the kinetics of lipogenesis [13]. Suitable temperature [13], salinity [19], aeration [38, 45], and mechanical stirring [43] have been reported to influence lipid production in *H. akashiwo*, however implementation of these lipid-regulating strategies, in certain cases, have resulted in decrease in total biomass [38, 43]. Notably, how hydrodynamic cues impact lipid production in *H. akashiwo*, and its collateral impact on biomass generation across physiological growth stages, have remained unknown. This opens up potential avenues to optimize algal lipid generation without compromising algal growth. Here, using two distinct strains of *H. akashiwo* we study how hydrodynamic cues, as a biophysical stressor [28], trigger lipid production in motile algal species under different scenarios. We quantify the lipid production concurrently with the biomass and photo-physiology. Specifically, our results reveal that the onset timing of the hydrodynamic cue, with respect to the growth curve, is a key determinant of enhanced lipogenesis and sustained fitness. Through rigorous monitoring of the growth kinetics, carrying capacity and photo-physiology, simultaneously with intracellular lipid production, we propose a hydrodynamic strategy for enhanced lipid production without compromising biomass as well as physiological fitness. This work presents hydrodynamic perturbation and its onset timing as tunable parameters to advance sustainable development of the next generation of microalgae-derived biofuels.

## II. MATERIALS AND METHODS

### A. Cell culturing in static conditions

Two strains of *H. akashiwo*, CCMP452 and CCMP3107 (hereafter referred to as HA452 and HA3107), were cultured under sterile conditions in 50 ml glass tubes using f/2-Si, a nutrient variant of the standard f/2 medium lacking silica. Artificial seawater (ASW), prepared in-house using 36 g of sea salt in deionized (DI) water, was infused with 1 ml of nitrates, 1 ml of phosphates, 1 ml of trace elements, and 0.5 ml of vitamins [13]. Three biological replicates for each strain were propagated from mother cultures in the exponential phase of their growth curve: 2 ml of cell culture was transferred from the upper part of the mother culture to 25 ml of fresh medium under sterile conditions under a laminar flow hood. The propagation process involved harvesting cells from the upper layer of the parent culture and transferring them into a nutrient-enriched medium for further growth. For this work, three biological replicates were used, each propagated in isolated culture tubes for at least 10 generations. For each strain, two sets of samples, each containing three culture tubes derived from the three mother cultures, were prepared. The cell cultures were incubated under a 14:10 h light/dark cycle to closely mimic natural diurnal rhythms, and maintained at 22°C under white light intensity of 1.35 mW during the light phases. The light phase commenced at 04:00 AM, and concluded at 06:00 PM to simulate day-night cycles. The light was automatically switched off from 06:00 PM to 04:00 AM of the following day to represent night-time conditions. All experiments were conducted between 10:00 AM and 04:00 PM, unless otherwise specified. Each strain was represented by three distinct biological replicates, with multiple technical replicates as outlined in dedicated sections of the text.

### B. Cell culturing under hydrodynamic perturbations

Cell cultures were prepared following the protocol outlined above. Thereafter, two different hydrodynamic configurations were investigated: *Scenario* 1 with 0-hour delay, wherein samples were exposed to hydrodynamic cues right after their inoculation in fresh media. The freshly inoculated cell culture were placed on an orbital shaker that was housed within the same incubator as the control culture. The shaker was operated at a speed of 110 revolutions per minute (rpm), introducing a hydrodynamic stimulus as the sole variable relative to the control sample. The orbital shaker provided a mild swirling action to stir the culture medium, thereby generating a hydrodynamic perturnation within the cell culture.

For the second configuration, i.e., *Scenario* 2 with 120-hour delay, the cell cultures underwent an initial static incubation period, similar to the control population. Once the cell cultures reached mid-exponential phase, they were subjected to the aforementioned hydrodynamic perturbations via an orbital shaker. This approach ensured that, aside from the introduced hydrodynamic cues, all samples experienced identical environmental factors ( see Fig. 1A), facilitating a controlled comparison of the effects of hydrodynamic versus static growth conditions.

**FIG. 1.**
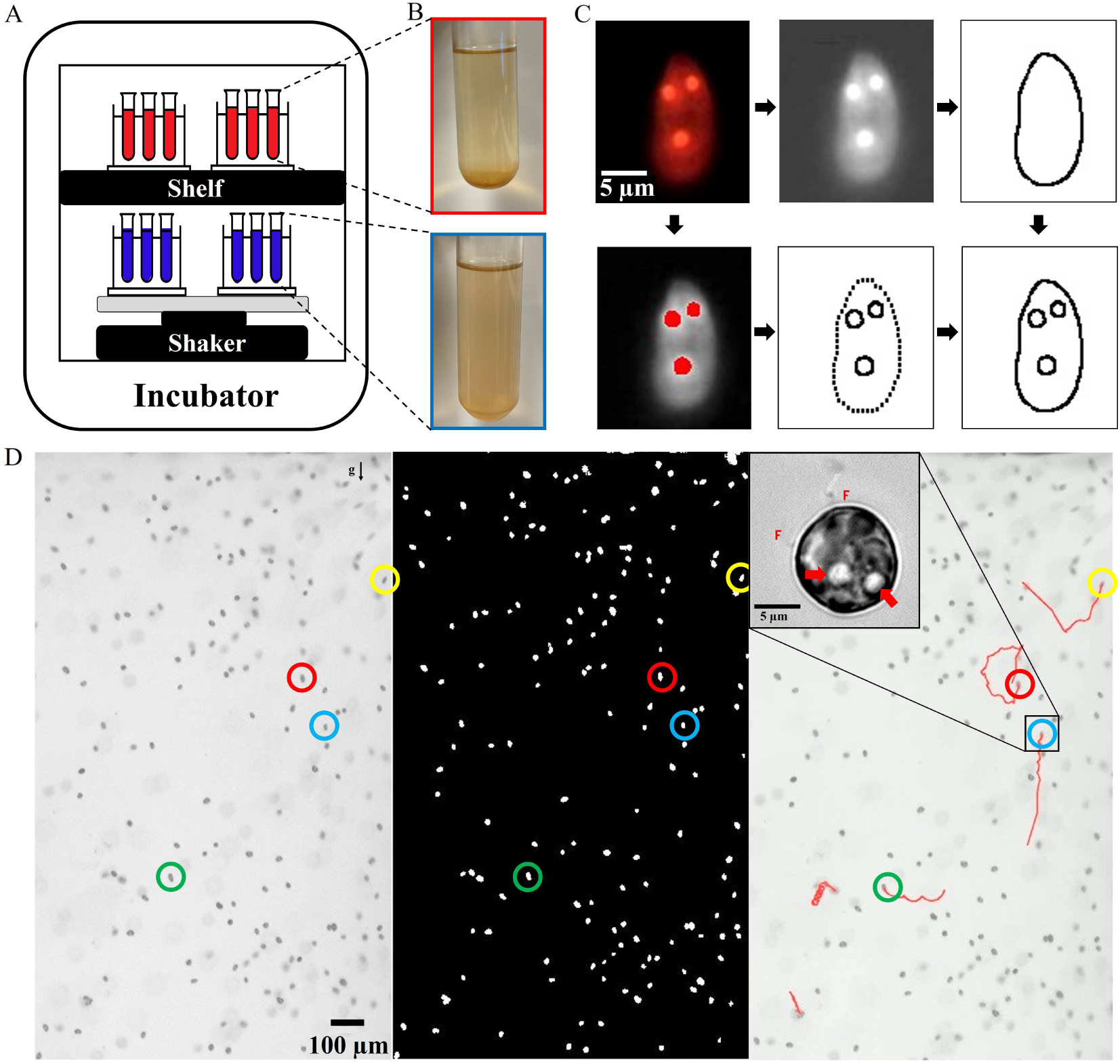
Experimental concept showing analysis pipeline for quantification of physiology and lipid generation. (A) The schematic illustrates our experimental concept: Two sets of cell cultures are housed within a temperature- and light-controlled incubator. The first set is maintained under static conditions (control population), while the second set is experienced hydrodynamic perturbation on an orbital shaker. The blue and red colour scheme is used throughout the article to indicate the populations on the shaker (exposed to hydrodynamic cues) and the static (control) populations respectively. (B) Snapshot of the two representative culture tubes, photographed here 180 h after inoculation. (C) Quantitative analysis of the intracellular lipids and the cell morphology, obtained using thresholding algorithms to estimate the relative volume of lipid droplets compared to the total cell volume. (D) Visualizing swimming cells in within a vertical millifluidic chamber allows quantification of the cell concentration and dynamics, with the direction of gravity indicated by the gravity vector ‘g’. The analyzed image (middle panel) displays bright spots representing cells after background subtraction. The right panel presents motile cells, with a few representative trajectories. The inset shows an *H. akashiwo* cell with its two flagella (indicated as F) and two lipid droplets appearing as bright spheres (indicated by two arrows), imaged at 100x magnification.

The hydrodynamic perturbation due to the orbital shaker was quantified following Gallardo Rodríguez *et al.* [46]. We calculated the energy dissipation rate once the system reached steady state condition (Eq. 1). The dissipation is calculated as the rate of energy dissipation per unit mass within the system throughout the period of perturbation, determined as follows:

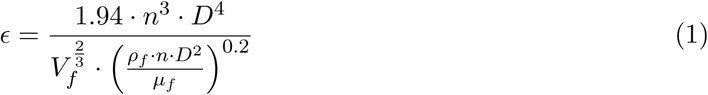

In the equation, *ɛ* represents the bulk mass-specific energy dissipation rate of system, where *n* signifies the rotational speed of the culture tube, while *D* refers to the tube’s diameter. The terms *V_l_*, *ρ_l_*, and *µ_l_* correspond to the fluid’s volume, density, and viscosity, respectively, with the fluid in this context being seawater. The volume of the fluid is 27 ml (25 ml of f/2-Si medium + 2 ml of mother cell’s suspension). The shaker was set at 110 rpm using, for which *ɛ* is calculated to be 8.78 *×* 10^−4^ W/kg in the system, which is an order of magnitude stronger than typical mild ocean turbulence [28].

### C. Variation of nutrient levels in microalgal cultures over time

To assess nutrient concentrations within the *H. akashiwo* culture, samples were collected at regular intervals throughout the growth period. The residual concentrations of nitrate and phosphate in the medium were quantified spectroscopically using a Spectroquant^®^ Prove 600 spectrophotometer, following the manufacturer’s protocols and reagent kits specifically designed for nitrate and phosphate analysis. Details of the protocol can be found in an earlier report [13].

### D. Quantifying *H. akashiwo* growth kinetics

To accurately assess cell concentration over time, small aliquots were collected from the culture tubes and counted using a microscope to establish a precise cell count. This was repeated for all experiments, spanning multiple biological and technical replicates. Starting 24 h after fresh incolutation, samples were drawn from the middle of the culture tube into a 3.4 *µ*m^3^ observation chamber. To ensure uniformity without imposing undue stress on the cells, each culture tube was gently stirred beforehand. Nikon^®^ SMZ1270 stereo microscope, capturing images at 16 frames per second (fps) over a span of 10 seconds was used to acquire dynamic images, which were then processed using MATLAB’s computer vision packages (Version 9.10.0.1669831) to count the cells in each frame, to finally arrive at the cell count. The aggregated cell concentration was plotted over time at intervals of 24, 48, 72, 96, 192, 264, 336, and 384 h, or until a plateau in cell concentration was reached, indicating the carrying capacity. Additionally, the cell count was validated by performing flow cytometry and particle tracking at selected time points chosen randomly. These independent validation confirmed the accuracy of our MATLAB code, with the results having a strong agreement across the methodologies employed (see Fig. 9).

The growth curve was then fitted to a logistic curve to determine the doubling time and carrying capacity of each population. When plotted in logarithmic scale, linear segments indicated the exponential growth phase, for which the doubling time (as well as the specific growth rates were extracted). The logistic growth model provides a quantitative approach to model population kinetics, modelled as:

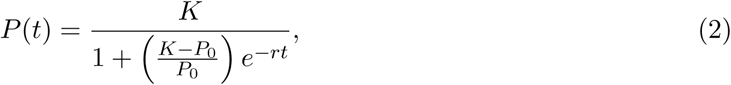

where *P* (*t*) is the population at any time *t*, *P*_0_ is the initial population size, *r* denotes the intrinsic growth rate, *K* is the carrying capacity, with *e* being the base of the natural logarithm. The doubling time (*T_d_*) can be derived as:

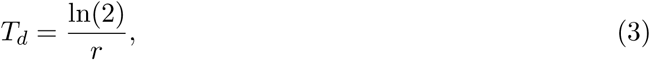

signifying that the *T_d_* is inversely related to the intrinsic growth rate *r*. The specific growth rate, *r*, captures the per capita rate of population increase under optimal conditions. Finally, the carrying capacity, *K*, signifies the maximum population size that the environment can sustain indefinitely, taking into account resource availability as well as biochemical constraints. It is estimated from the plateau phase of the growth curve. Estimating the parameters *r* and *K* from empirical data involves nonlinear regression, fitting the logistic growth equation to observed population sizes over time (see Fig. 8). This allows us to quantify the doubling time during the exponential growth phase and provides measureable insights into the long-term population health under the evolving environmental constraints.

### E. Flow cytometry

To ensure accurate cell concentration measurements and validate the cell counting code, flow cytometry was used to quantify cell concentration in randomly selected samples, enabling iterative optimization of the code. Through this process, the discrepancy between the cell counting code and flow cytometry results was reduced to less than 10%. Flow cytometry was conducted using an *Attune™NxT Acoustic Focusing Cytometer* in volumetric mode, which facilitated direct cell counting without the use of reference beads. Samples with estimated concentrations exceeding 10,000 cells (post 100–120 h of inoculation) were diluted in Milli-Q® water to prevent inaccuracies due to high concentrations. Algal cells were identified based on intrinsic chlorophyll autofluorescence captured by the BL3-H channel. Sequential gating strategies were applied, beginning with fluorescence-based gating to isolate autofluorescent cells, followed by size and complexity assessments using forward scatter (FSC) and side scatter (SSC) plots (see Fig. 9). Final cell concentrations were calculated from the total detected events, adjusted for dilution factors and sample acquisition volumes, and were further compared to results from image-based cell analysis scripts for validation.

### F. Measuring lipid production using cell-scale microscopy

Biosynthesis and accumulation of intracellular lipid droplets (LDs) were conducted at predetermined intervals. The cell cultures were gently homogenized to ensure uniform cell density. Using Nile Red (Thermo Fisher Scientific, excitation/emission at 552/636 nm), a fluorescent dye specific for neutral lipids, was used to stain the LDs. 10 *µ*l of 100 *µ*M Nile Red in dimethyl sulfoxide (DMSO) was dissolved in 200 *µ*l of *H. akashiwo* culture supernatant. This mixture was then subjected to vigorous vortexing for even dispersion. Subsequently, 200 *µ*l of the cell suspension was introduced to the Nile Red solution to achieve a final dye concentration of 2.4 *µ*M, followed by a 15-minute incubation in complete darkness at room temperature. We combine phase contrast and fluorescence microscopy to acquire the LD accumulation in individual cells, using an Olympus CKX53 inverted microscope equipped with a high-resolution color camera (Imaging Source, DFK33UX265). To mitigate any phototoxic effects during data acquisition, the intensity of the excitation LED (552 nm) was carefully maintained at a low setting, with a peak intensity of 5 on the relative scale provided.

We measure the cell area, LD dimensions (area and volume), and their spatial distribution within the cells from the image datasets obtained as a time series of 16 fps for a duration of about 10 seconds. A total of 20 cells were studied in each case, representative images for the two scenarios (Scenarios 1 and 2 described above) are shown in Figs. 2 and 3. The LD dimensions was obtained from their contour areas, which were obtained by image thresholding techniques using custom-built MATLAB image processing scripts alongside ImageJ software. Following analysis, the maximum and minimum Feret diameters of each LD were extracted through threshold-based segmentation. The Feret diameters were then used to estimate the volume of each LD, assuming a prolate spheroid geometry. Subsequently, the radius of an equivalent sphere, possessing a volume equal to that of the calculated LD, was determined. This equivalent radius was utilized to estimate the cross-sectional area of the LD, providing a consistent and standardized metric for LD comparison across different samples. Additional details on the LD quantification can be found in Ref. [13]. The same procedure was then applied to the cell itself wherein we determined its cross-sectional area (see Supplementary Figure 10). The normalized lipid size was obtained as the ratio of the lipid area to the cell area, by dividing the cross-sectional area of the effective lipid droplet by that of the corresponding cell.

**FIG. 2.**
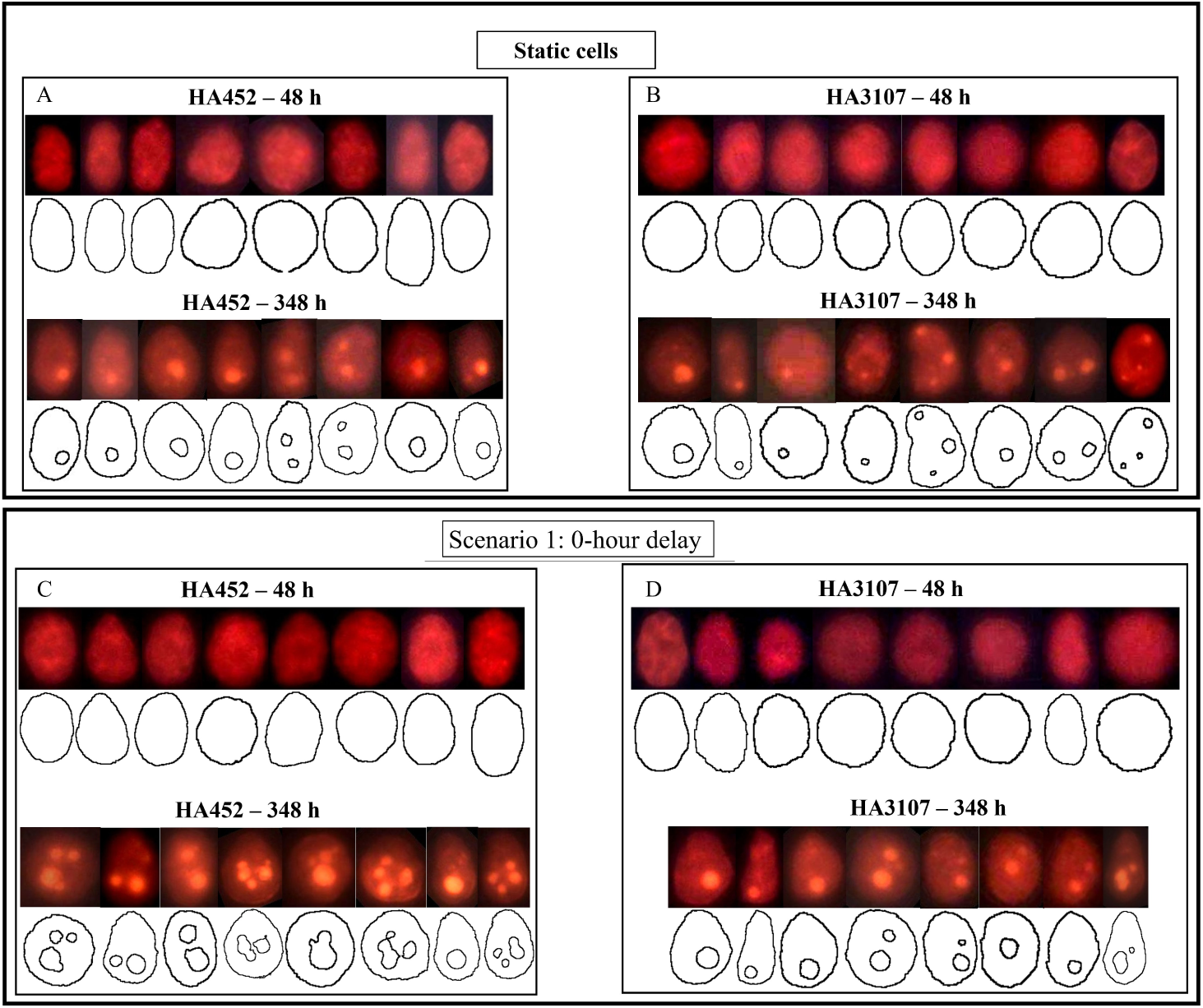
Scenario 1 (0-hour delay): Quantifying lipid production in *Heterosigma akashiwo* using single cell fluorescence microscopy. (A)–(D): Representative fluorescent images, which were analysed to obtain the lipid droplet dimensions and cell contours for both strains of *H. a akashiwo*, shown here for the Scenario 1 (0-hour delay), for early exponential (48 h) and stationary growth phases (348 h). The neutral LDs appear bright orange. Control (static) populations are shown in panels (A) and (B) for HA452 and HA3107 respectively, while corresponding hydrodynamically perturbed populations are shown in panels (C) and (D). Under static conditions, the average lipid accumulation increased from 1.0 *±* 0.5 *µ*m^3^*/*cell at 48 h to 22.0 *±* 18.1 *µ*m^3^*/*cell at 348 h (*n* = 60 cells) for HA452; and from 2.0 *±* 3.0 *µ*m^3^*/*cell at 48 h to 15.2 *±* 12.0 *µ*m^3^*/*cell at 348 h (*n* = 60 cells) for HA3107. (C) HA452 population subjected to 0-hour delay perturbation at 48 h (top row) and 348 h (bottom row), reveal an increase in cytoplasmic lipid accumulation from 5.0 *±* 12.0 *µ*m^3^*/*cell at 48 h to 170.3 *±* 58.6 *µ*m^3^*/*cell at 348 h (*n* = 60 cells). (D) HA3107 under 0-hour delay perturbation showed an increase in cytoplasmic lipid accumulation from 4.2 *±* 2.0 *µ*m^3^*/*cell at 48 h to 49.1 *±* 31.5 *µ*m^3^*/*cell at 348 h (*n* = 60 cells).

**FIG. 3.**
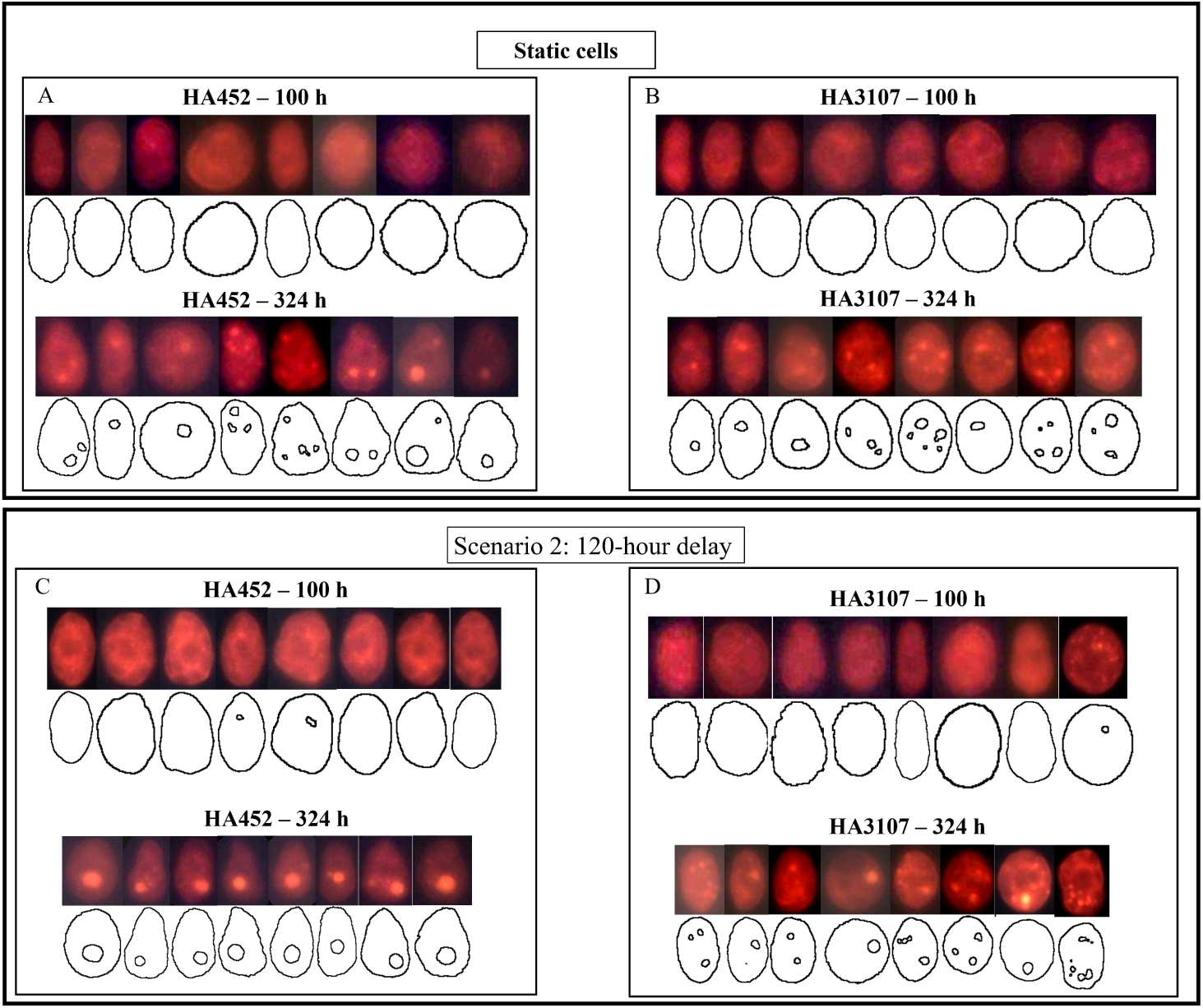
Scenario 2 (120-hour delay): Quantifying lipid production in *H. akashiwo* using single cell fluorescence microscopy. (A)–(D): Representative fluorescent images, which were analysed to obtain the lipid droplet dimensions and cell contours for both strains of *H. a akashiwo*, shown here for the Scenario 2 (120-hour delay), for exponential (120 h) and stationary growth phases (324 h). The neutral LDs appear bright orange. Control (static) populations are shown in panels (A) and (B) for HA452 and HA3107 respectively, while corresponding hydrodynamically perturbed populations are shown in panels (C) and (D). The average lipid accumulation increased from 1.4 *±* 0.8 *µ*m^3^*/*cell at 100 h to 21.6 *±* 4.1 *µ*m^3^*/*cell at 324 h (*n* = 60 cells) for static HA452 populations. Correspondingly, lipid accumulation varied from 6.2*±*0.4 *µ*m^3^*/*cell at 100 h to 12.9*±*4.5 *µ*m^3^*/*cell at 324 h (*n* = 60 cells) for HA3107 under static condition. (C) Upon exposure to hydrodynamics cues after a 120-hour delay, HA452 cells had accumulated 5.8*±*2.6 *µ*m^3^*/*cell at 100 h, which significantly increased to 48.2 *±* 3.5 *µ*m^3^*/*cell at 324 h (*n* = 60 cells). (D) Lipid accumulation in HA3107 cells subjected to a 120-hour delay perturbation increased marginally from 6.9 *±* 1.6 *µ*m^3^*/*cell at 100 h to 16.2 *±* 1.3 *µ*m^3^*/*cell at 324 h (*n* = 60 cells).

### G. Evaluating changes in photophysiology due to hydrodynamic cues

Key photosynthetic indices, including the maximum quantum yield of Photosystem II (PSII), *F_v_/F_m_* (hereafter referred to as photosynthetic efficiency), maximum electron transport rate (*ETR_max_*), and non-photochemical quenching (*NPQ*) were measured using a Multi-Color-PAM Chlorophyll Fluorometer (Heinz Walz GmbH, Effeltrich, Germany). 1.5 ml aliquot of each population was was carefully extracted from each culture tube, transferred into quartz-silica cuvettes (Hellma absorption cuvettes; spectral range: 200–2500 nm; path length: 10 mm), and dark-adapted for 5 minutes. Longer dark adaptation durations did not impact the results, as reported in detail elsewhere [13]. *F_v_/F_m_*, *ETR_max_*, and *NPQ* values were calculated using the PAM-Win software (accompanying the PAM fluorometer), under both saturation pulse (SP) technique as well as the light curve methods. This ensures a comprehensive analysis of photosynthetic performance. The *NPQ* value at 500 photosynthetically active radiation (PAR) was specifically interpolated from the *NPQ*-PAR series provided by the software, allowing for precise characterization of the non-photochemical quenching response under this light intensity.

### H. Statistical tests

We conducted paired t-tests to compare the total lipid accumulation, normalized lipid droplet area, and photophysiology for the control populations (no perturbations) as well as the experimental populations (both Scenarios 1 and 2) at regular intervals on the growth curve, spanning early exponential to the stationary growth stages. For the statistical analysis of the growth kinetics (carrying capacities and doubling times) between the control and experimental groups, we utilized two-sample t-tests with the significance level set at *P* = 0.05. All statistical analyses were performed using GraphPad Prism software.

## III. RESULTS

### A. Hydrodynamic cues expedite growth but maintain carrying capacity

We quantify and compare the growth kinetics of freshly inoculated populations exposed to hydrodynamic cues (Scenario 1) with their control counterparts. Using microscopy and image analysis as detailed in Section II D (Fig. 1D), we computed the cell concentration as a function of time. Figure 4A and D present the growth curves for HA452 and HA3107 populations for both the control and hydrodynamically perturbed cases. The shaded region around the solid-line indicates the standard deviation, while the solid line represents the mean value measured across all replicates. We fit a logistic function (bottom right inset panels for each growth curve) to derive their doubling time (Supplementary Figure 8) and the carrying capacity. For HA452, the doubling time drops significantly (Fig. 4B), while the carrying capacity remains relatively unchanged (Fig. 4C). This indicates that the population exposed to hydrodynamic cues achieve their maximum concentration more rapidly. Mechanical agitation right after inoculation does not inhibit normal biomass yield but significantly decreases the time required to reach maximum biomass, thereby increasing the specific growth rate.

**FIG. 4.**
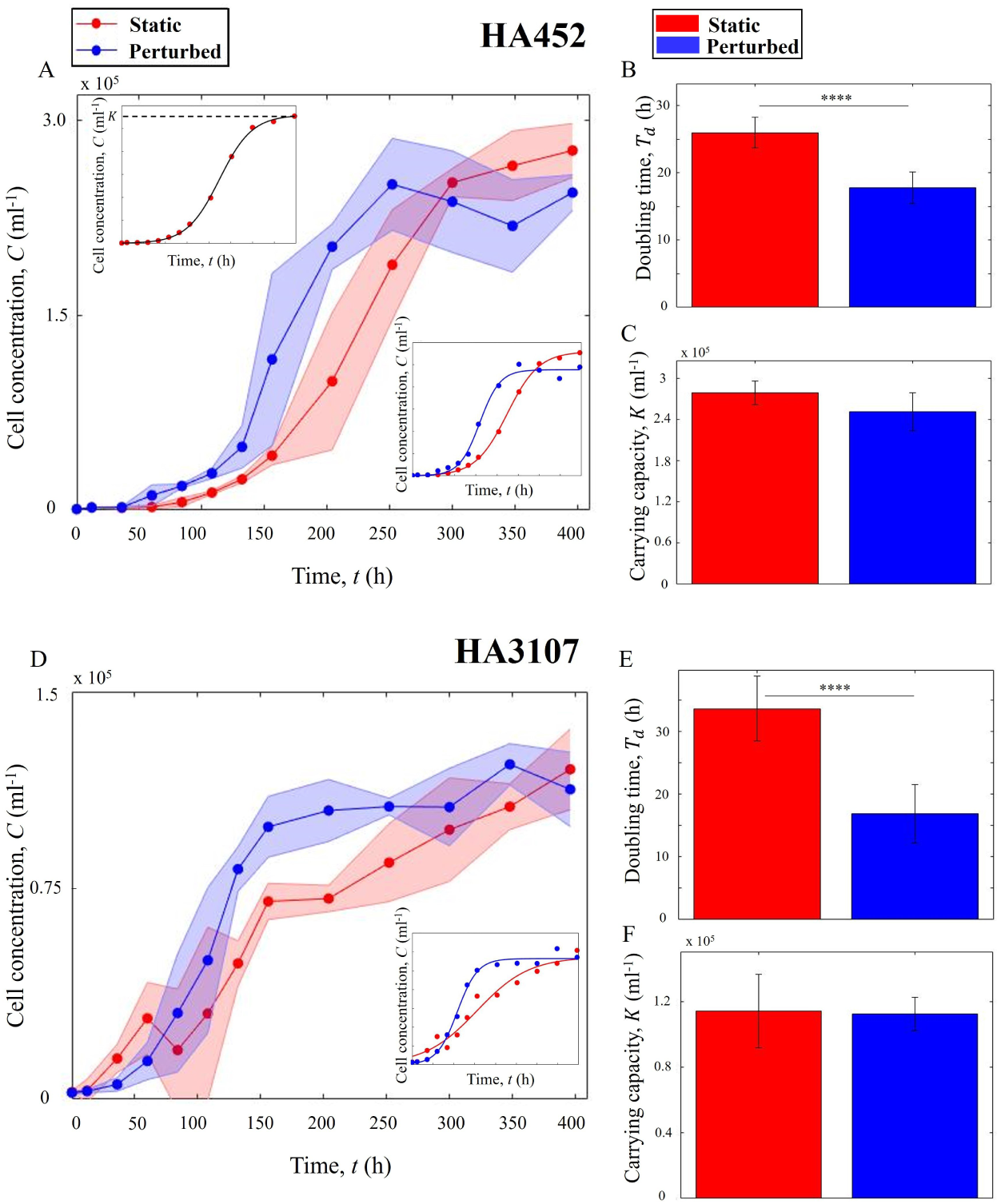
Hydrodynamic cues promote algal growth without impacting population size. (A) Comparative growth curves of HA452 under static (red) and hydrodynamic cues (blue). The plot shows the average cell count across all biological and technical replicates. The solid line connects the mean values, while the shaded area indicates the standard deviation. Upper inset shows a fitted logistic curve on sample growth data, where *K* indicates the carrying capacity. The lower inset presents growth curves fitted by a logistic function for static and perturbed cells. (B) Comparative doubling times for the two populations in (A). The bar plots show the mean doubling time for each subset, with error bars representing the standard deviation. The data indicates that hydrodynamic perturbation immediately after inoculation significantly reduces the doubling time (*P <*0.0001), accelerating HA452 cell growth. (C) Estimated carrying capacities for the two populations in (A), show that the maximum population size remain comparable across both conditions (*P >*0.05). (D) Growth curves of HA3107 cultures under static and hydrodynamic cues, the inset plots refer to similar plots as described above. (E) Doubling times for the two populations in (D) indicate that the hydrodynamic cues significantly reduces the doubling time (*P <*0.01), while the carrying capacities (F) show minimal change (*P >*0.05). For each data point, samples were obtained from three different biological replicates, each sampled at least twice (yielding two technical replicates).

For HA3107 population (Fig. 4E and 4F), a similar trend is observed, i.e., the population reached the carrying capacity significantly earlier than the control population. Statistical analysis indicates that the maximum biomass yield did not differ significantly between the control and perturbed cells (Fig. 4F). However, the specific growth rate increased in the perturbed cells by over 100%, resulting in a significantly reduced doubling time (Fig. 4E). These results confirm that the initiation of hydrodynamic perturbation shortly after fresh inoculation (within a couple of hours) is a potent method to promote faster growth without impacting the population fitness.

### B. Onset timing of the hydrodynamic cues tunes the maximum population size

In Scenario 2 of our studies, we investigate the impact of the onset timing of hydrodynamic cues on the growth kinetics and population size. The cell cultures were initially maintained under static conditions, and upon reaching the mid-exponential growth phase (*≈* 120 h after inoculation), they were exposed to perturbations. This specific time point was selected based on the populations’ physiology, as observed in our current experiments as well as previously reported growth kinetics for HA452 [28] and HA3107 [13]. Subsequent steps to quantify the growth curve were similar to those outlined in sections II and III A.

As shown in Figure 5, the carrying capacity varied significantly between the control and the perturbed populations for both strains (panels C and F of Fig. 5). On the other hand, the doubling times remained unaffected, as depicted in Figures 5B and 5E respectively for HA452 and HA3107. This is in contrast to the Scenario 1 (cells were subjected to perturbation immediately after inoculation, i.e., 0-hour delay), where the carrying capacity remained stable while the doubling time dropped significantly (Fig. 4). Conversely, in the Scenario 2, a significant reduction in carrying capacity is noted, with non significant variation in the doubling times.

**FIG. 5.**
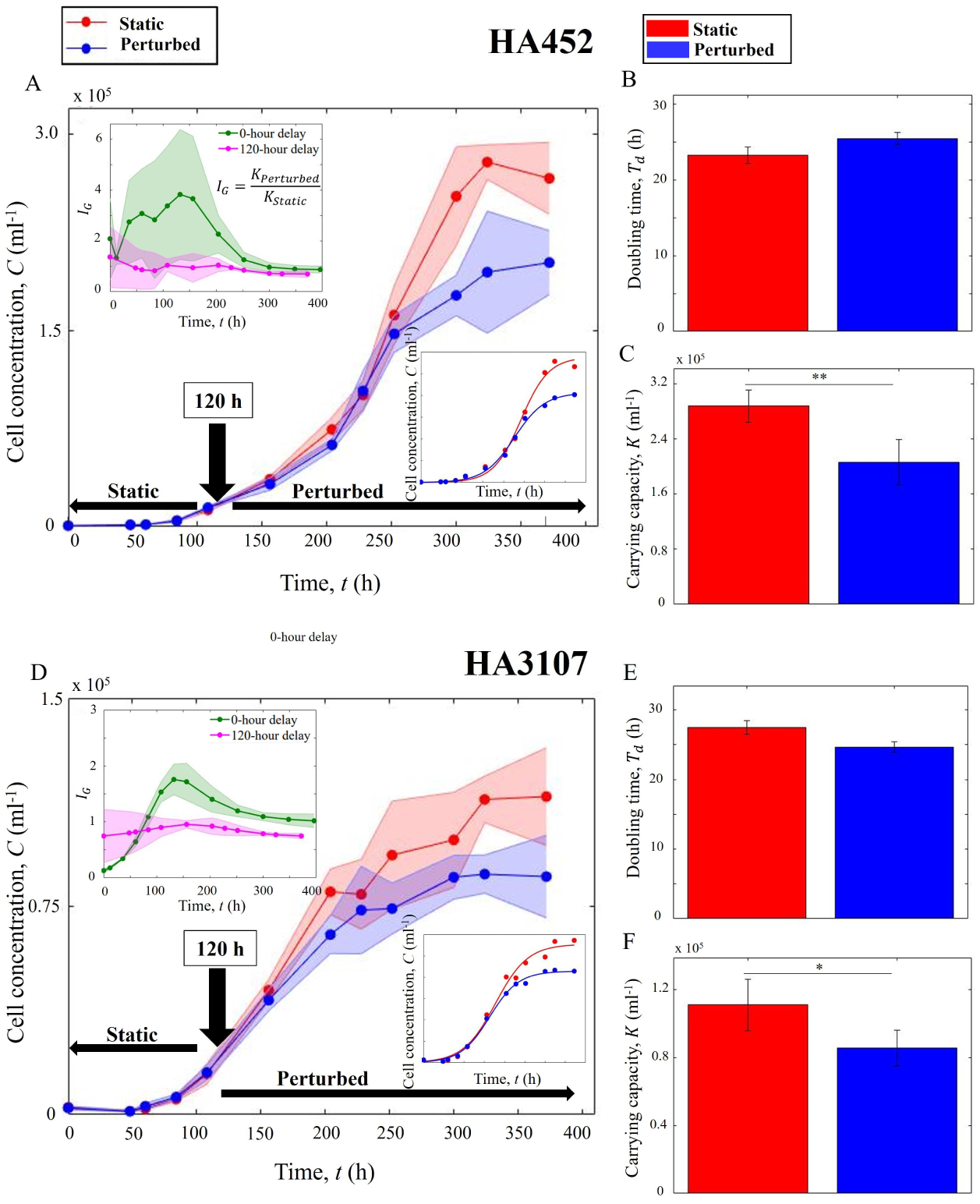
The onset timing of hydrodynamic cues tunes the population size. (A) Growth curves of HA452 cultures under Scenario 2 (120-hour delay, shown in blue) relative to the control population (red). The transition time point at which the hydrodynamic cues are introduced is shown by the black arrow. Upper inset compares the growth indices for Scenario 1 (0-hour delay, green) versus Scenario 2 (120-hour delay, pink). The lower inset shows the logistic-fitting on the cell concentration. (B) No significant difference in the doubling times for the two scenarios is noted (*P >* 0.05). (C) The maximum population size, i.e., the carrying capacity, shown as bar plots (error bars indicate standard deviation), is significantly suppressed in case of the perturbed sample (blue) relative to the control population (red), *P <* 0.01. (D) Growth curves for two subsets of HA3107 cultures, grown under the same conditions as HA452 cells, follow a similar trend. The upper inset illustrates the growth indices, while the lower inset displays the logistic fitting for the cell concentrations. (E) Hydrodynamic cues introduced at the mid-exponential growth phase does not impact the doubling time (*P >* 0.05), however the carrying capacities for the two sets of populations differ significantly (*P <* 0.05). For each data point, samples were obtained from three different biological replicates, with each sample having two technical replicates.

We define a growth index, *I_G_*, to quantify the relative change in population size for both the scenarios spanning the growth phases (upper inset in Figures 5A and D. The green plot indicates the 0-hour case, while magenta curve plots the same for the 120-hour delay case. The growth index, denoted by *I_G_*, is defined as:

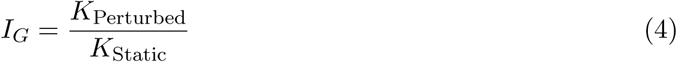

where *K*_Static_ represents the cell concentration of the static cells at each time point, derived from the corresponding logistic model, and *K*_Perturbed_ represents the cell concentration of the perturbed cells at each time point, also derived from their corresponding logistic model. During the initial phase, the negative slope for the 0-hour delay plot is steeper than for the 120-hour delay plot, indicating that cells under immediate perturbation have limited ability to grow, produce biomass and adapt. Consequently, these cells produced less new biomass compared to the control cells. After approximately one day, the slope of the plot remained positive up to around 150 h. This indicates that cells subjected to hydrodynamic perturbation immediately after inoculation grew favourably and produced more biomass until reaching their mid-exponential phase. Due to their rapid growth, even when the slope of their growth index became negative again after 150 h, their cell concentration remained higher than that of the static cells until around 300 h. The growth index for the 0-hour delay perturbation remained higher than 1 throughout the stationary phase, decreasing to approximately 0.85 after 400 h (late stationary phase). In case of Scenario 2 (120-hour delay), *I_G_* fluctuated around 1 before the transition point. When the cells were subjected to hydrodynamic perturbation, *I_G_* remained marginally above 1 for around 100 h, potentially due to homogenized nutrient distribution, which facilitated higher nutrient access for the cells [38]. Subsequently, the index reduced to less than 1, eventually decreasing to approximately 0.69.

Taken together, these results indicate that algal populations, when exposed to hydrodynamic cues early on in their growth phase, proliferated at a faster rate, achieving the same carrying capacity within a shorter time. In contrast, delayed exposure to hydrodynamic cues allowed the population to grow at comparable rates during the exponential phase but transitioned to their stationary phase sooner than the control samples. Thus, freshly inoculated motile cells grown under hydrodynamic perturbations increased their productivity relative to the control cells grown under static conditions. Comparing *I_G_* values for HA452 and HA3107 for the 0-hour delay scenario, a longer adaptation period is noted for HA3107 when exposed to hydrodynamic perturbations. From around 96 h to 150 h (the exponential growth phase), the perturbed cells exhibited higher biomass production while continuously subjected to hydrodynamic perturbation. After 150 h, the slope turned negative, indicating a higher growth rate for the control cells compared to the perturbed ones. Ultimately, this resulted in equivalent carrying capacities for both subsets of cells in the 0-hour delay scenario.

For the Scenario 2, the growth index exhibited an immediate increase following the introduction of the perturbation, reaching its maximum value. This increase is likely attributable to a more uniform nutrient distribution within the cultures of the perturbed populations. Previous studies have demonstrated that cellular stress may cause HA3107 to exhibit diffusive than ballistic behavior [13], which can further restrict their access to nutrients. The growth was subsequently suppressed due to hydromechanical forces, stabilizing at around 0.74. This stabilized value is higher than the corresponding index for HA452 cells, indicating that HA3107 cells are less adversely affected by the 120-hour delay perturbation scenario. The onset timing of hydrodynamic cues thus offers a novel handle to regulate algal growth kinetics. While the extent of these responses are strain-dependent, the observed trends were found to be similar for both the growth rate as well as the carrying capacity.

### C. Algal lipid production depends on the onset timing of hydrodynamic cues

Lipid accumulation was systematically monitored throughout both experimental scenarios (0-hour delay and 120-hour delay scenarios). Cell samples were collected at regular intervals from all culture tubes, in accordance with the protocols outlined in section II F. Unlike previous studies [38, 45, 47], we have quantified lipid accumulation at the single cell level using epifluorescent microscopy (Fig.1C), with additional validation using bright-field imaging at 100X magnification (Fig. 1D, inset). For the 0-hour delay scenario (Fig. 6A), the normalized lipid area was consistently higher for the populations exposed to the hydrodynamic cues, indicating a sustained enhancement of the lipid production across all time points (while the cell area remained constant, Supplementary Fig. 10A)). Despite similar biomass yields between the control and perturbed cells, as shown in Figure 4C, the normalized lipid area in perturbed cells increased notably during the stationary phase, resulting in an almost 4-fold enhancement upon exposure to hydrodynamic cues (*P <* 0.01) after 350 h. While nutrient limitation affects both the control and perturbed cell cultures by the time they reach the early stationary phases [13], nutrient stress alone cannot account for the significantly higher lipid accumulation observed in the populations exposed to hydrodynamic cues. Depletion of nitrates and phosphates (see Supplementary Figure 12) are known to induces cellular stress, leading to increased lipid production and accumulation [13], our results indicate that hydrodynamic cues allow significantly higher enhancement beyond the nutrient-limited lipogenesis.

**FIG. 6.**
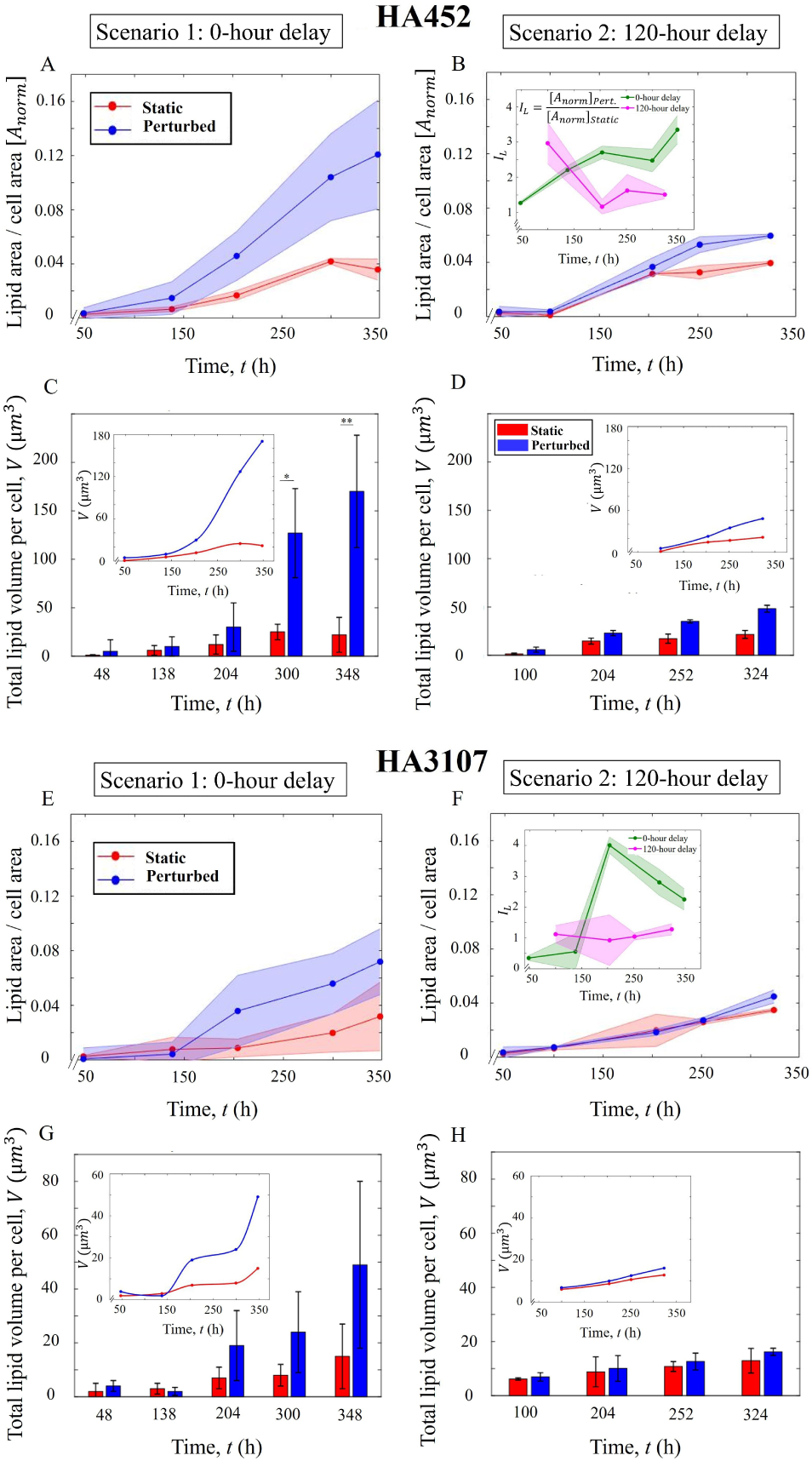
Onset timing of hydrodynamic cues modulates enhancement of lipid production. (A) Normalized lipid area across the growth phases under Scenario 1 (0-hour delay), for HA452. Solid curve represents the average ratio over the replicates, while the shaded area shows the standard deviation. Significant increment of lipid production (*>* 3-fold, *P <* 0.001) is noted. (B) Delayed exposure to hydrodynamic cues (120-hour delay) results in a less pronounced increment of lipid (*P <* 0.05). The inset shows the index of lipid accumulation, 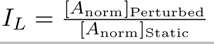 for both scenarios across the growth phase. (C) Total lipid volume per cell shows a statistically significant increase compared to the control cells (*P <* 0.01) during the stationary phase. The inset captures the trend during the course of experiment. (D) Total lipid volume per cell for the 120-hour delay scenario exhibited lower increment relative to the 0-hour delay case. The inset shows the corresponding trends during the course of the experiment. (E) Normalized lipid area for HA3107 under Scenario 1 (0-hour delay) shows a marginal increment (*p* = 0.555), while for Scenario 2 (F), an increase in the normalized lipid area was noted, which was lower compared to the 0-hour delay scenario. The inset shows the variation of *I_L_* which was around 2.25 at the end, indicating larger lipid droplets for the perturbed cells. For Scenario 2, *I_L_* remained *≈*1 throughout, with a slight increase during the stationary growth stage for both control and perturbed cells. (G) Total lipid volume per cell for HA3107 cells under 0-hour delay scenario showed no significant enhancement (*p* = 0.0745). The inset shows the trend over the entire duration of the experiment. (H) The total lipid volume per cell in HA3107 under the 120-hour delay condition showed marginal enhancement.

For the Scenario 2 (120-hour delay, Fig. 6B), the normalized lipid area remained comparable between control and perturbed cells for approximately 100 h following the onset time point. However, upon reaching the stationary growth phase, a significant increase in the normalized lipid area was observed for the perturbed cells compared to the control cells (*P <* 0.05). This increase in lipid accumulation was less pronounced than in Scenario 1: an increase of 150% compared to an increase of *≈*400% observed in the 0-hour delay scenario. To quantify the lipid production trends, we define an index of lipid accumulation, *I_L_* for each scenario (Fig. 6B inset panel) as follows:

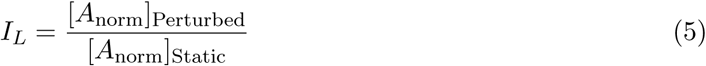

Here [*A*_norm_]_Perturbed_ and [*A*_norm_]_Static_ represent the mean normalized lipid area for the perturbed and static populations respectively. In the 0-hour delay case, *I_L_* showed a steady increment, with the steeper slope observed during the exponential phase. As demonstrated in section III A, HA452 cells in their exponential phase under hydrodynamic perturbation displayed accelerated growth and a significantly reduced doubling time compared to static cells. The lipid index indicates that these cells while growing at a faster rate than static cells, were concurrently accumulating more neutral lipids. The formation and storage of lipid droplets are biomarkers of cellular stress [13], which is known to suppress growth [45]. However, our results suggest an adaptive strategy whereby the HA452 population subjected to a stressful environment from the outset, allowing them to thrive and accumulate lipids while sustaining a steady fitness.

As the population entered the early stationary phase, the *I_L_* stabilised due to a drop in the cell concentration of the perturbed population relative to the static population. It then sharply increased starting from the mid-stationary phase, where a combination of hydrodynamic and nutrient-limitation induced cellular stress in both populations [13]. For the 120-hour delay scenario, the lipid accumulation index exceeded 1 prior to the hydrodynamic cues, with both populations initially exposed to identical environmental conditions. Despite introduction of hydrodynamic cues 120 h post-inoculation, *I_L_* continued to drop for 100 h post-perturbation, likely a more homogeneous nutrient distribution (due to shaking), thereby offering temporary adjustment to the hydrodynamic perturbation. *I_L_* eventually stabilized at around 1 after 100 h post-perturbation, as also reflected in the trends observed for the growth index (inset, Fig. 5A) and the normalized lipid area (Fig. 5B), suggesting that the cells were able to maintain their physiological state for *≈*100 h following exposure to the hydrodynamic stressor. Overall, the results indicate that algal populations exposed to hydrodynamic cues freshly after inoculation, are not only able to successfully maintain their physiological fitness, they also yield higher lipid volumes. Conversely, hydrodynamic cues applied on populations grown under static conditions for some time (here, until mid-exponential phase) led to reduced biomass alongside significantly lower lipid accumulation.

For HA3107 cells, the normalized lipid area for both control and perturbed cells is depicted in Figure 6E for the 0-hour delay scenario and in Figure 6F for the 120-hour delay scenario. For the Scenario 1, lipid accumulation increased more than that for the control populations, at around 140 h after inoculation. From this time point onward, the accumulated lipid in the perturbed cells remained consistently higher than in the control cells. Although this parameter more than doubled in the perturbed cells compared to the control ones, we could not establish statistically significant difference. Therefore, for HA3107 cells under the 0-hour delay scenario, the cells grew at a higher rate with a notably reduced doubling time, resulting in a similar biomass over a shorter timescale, with more than double the amount of accumulated lipid. For Scenario 2 (Fig. 6F), increase in the lipid accumulation did not reach the corresponding value observed for the 0-hour delay scenario. Taken together, reduction of the delay time (i.e., an early onset time for hydrodynamic cues) produced more cells with a higher amount of algal lipids, providing a tractable parameter space to enhance algal lipid productivity. Furthermore, the role of hydrodynamic cues on the onset and volume of lipid accumulation is strain-dependent. As presented in Figures 2 and 3, a markedly higher accumulation of lipid droplets is observed for HA452 relative to HA3107, particularly in populations which were exposed to hydrodynamic cues right after inoculation. The strain-specific difference in the accumulated lipids was less pronounced when the onset time for hydrodynamic perturbations was delayed.

### D. Onset timing of hydrodynamic perturbation impacts algal photo-physiology

Hydrodynamic cues impact photo-physiology [27], yet the role of the onset timing on photophysiology and possible induction of cellular stress remains unexplored. We quantify the maximum quantum yield of Photosystem II (PSII) photochemistry, represented by the ratio of maximum variable fluorescence (*F_v_*) to maximum fluorescence (*F_m_*) of chlorophyll (*F_v_/F_m_*) [13]. This ratio, or maximum quantum yield, has been extensively utilized in numerous studies as a standard metric for photosynthetic efficiency and indicator of cellular fitness [48]. Using a Multi-Color Pulse Amplitude Modulation (PAM) Chlorophyll Fluorometer (see Materials and Methods, Sec. II), we carry out a time series measurement of photo-physiology for the control and perturbed populations under both hydrodynamic scenarios described previously.

Figures 7A and 7B present the photosynthetic efficiency of HA452 under the 0-hour and 120-hour delay scenarios, respectively. Populations exposed to hydrodynamic cues immediately post-inoculation (0-hour delay) yield comparable photosynthetic efficiencies up to 200 h (Fig. 7A), and thereafter showed significant decline in performance spanning the stationary growth phase. On the other hand, the corresponding *ETR*_max_ (see Supplementary Fig. 11) remained within the range observed for the control populations. This constancy suggests that, for at least the initial 200 h, the cells neither compensated for the hydrodnamic perturbation nor responded to increased energy demands to maintain their photosynthetic performance [49]. Consequently, their photo-physiological status is considered stable. However, post the 200-hour mark, both the maximum quantum yield of Photosystem II and *ETR*_max_ exhibited declines, indicating cellular stress or damage to the photosynthetic apparatus. Notably, an elevation in *ETR*_max_ could potentially serve as a compensatory mechanism to sustain photosynthetic performance despite reduced PSII efficiency, a response that was absent in this scenario.

**FIG. 7.**
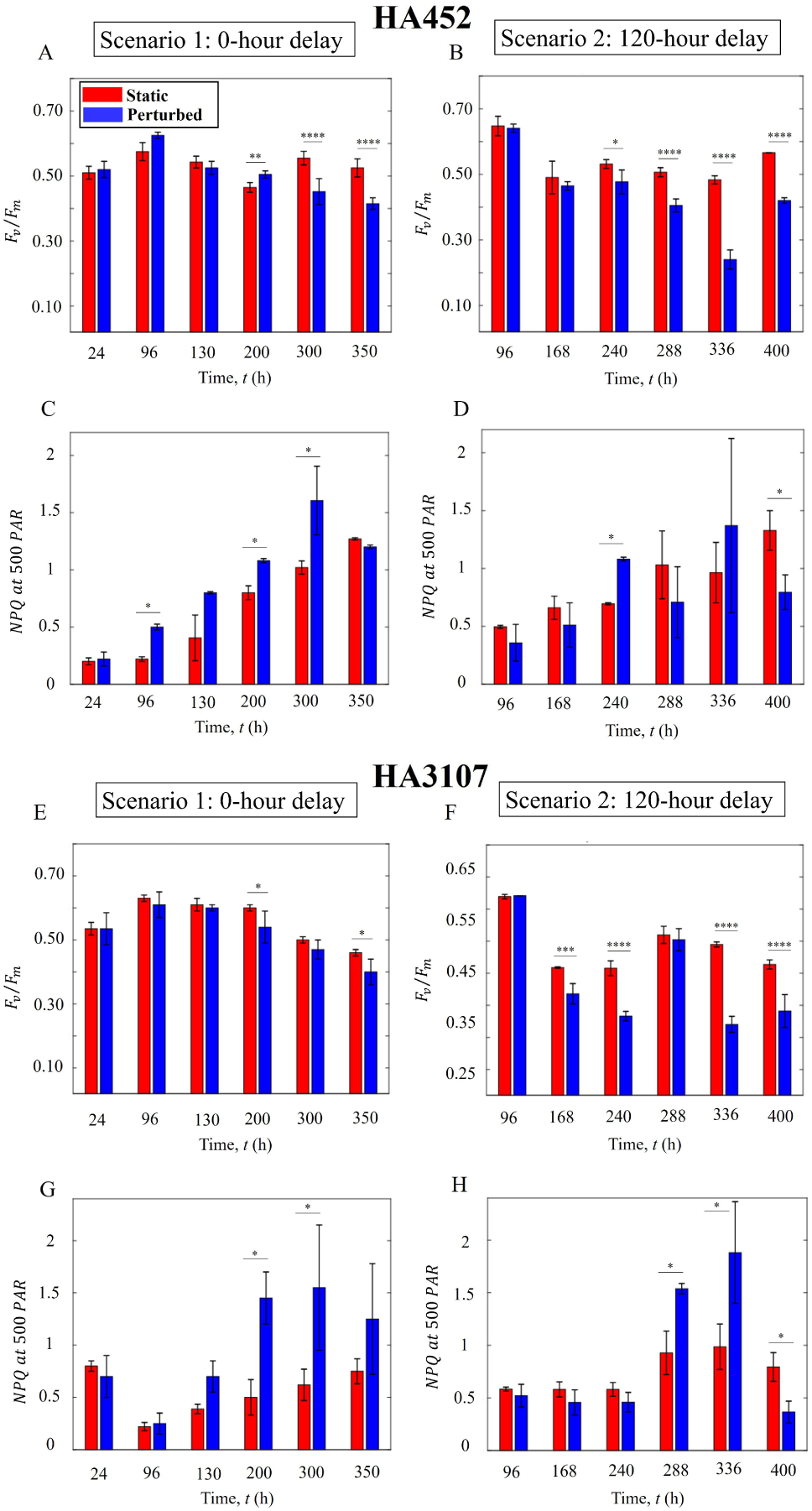
Onset timing of hydrodynamic perturbation impacts algal physiology. (A) Photosynthetic performance of HA452 under Scenarion 1: Perturbed cells exhibited comparable photosynthetic activity relative to static cells for up to 11 days, thereafter dropping significantly (*P <* 0.001) around 300 h post-inoculation. (B) Scenario 2: Following the initiation of hydrodynamic cues, a significant decrease in photosynthesis was observed within 5 days (*P <* 0.001). (C) HA452 *NPQ* values show comparable values initially, which later increased siginificantly (*P <* 0.05), eventually reaching a plateau before decreasing. (D) *NPQ* values of HA452 cells under the 120-hour delay scenario. (E) Photosynthetic efficiency of HA3107 cells under the 0-hour delay experimental condition. The perturbed cells maintained comparable photosynthetic efficiencies to the control cells throughout the experiment until a significant reduction was noted during stationary phase. (*P <* 0.05). (F) Photosynthetic efficiency of HA3107 cells under the 120-hour delay experimental condition: photosynthetic efficiency was significantly affected, with a marked decline observed at 168 h (*P <* 0.001). (G) HA3107 *NPQ* values under the 0-hour delay scenario, and under the 120-hour delay scenario (H). The *NPQ* value never increased in the perturbed cells, while *F_v_/F_m_* and *ETR*_max_ were significantly reduced, indicating their inability to self-regulate photophysiological responses to the hydrodynamic perturbations.

The non-photochemical chlorophyll fluorescence quenching (*NPQ*), an indicator of the excess absorbed light energy that is ultimately dissipated after conversion into heat [50], provides insights into the protective mechanisms against light-induced stress [51]. By interpolating the *NPQ* at 500 *µ*mol*/*(m^2^*/*s) akin to the light intensity typically observed at the ocean surface [13], we evaluate the *NPQ* values across all samples utilizing the series of *NPQ* values as a function of *PAR* recorded by the PAM fluorometer (Materials and Methods, Sec. II). As depicted in Figure 7C, freshly inoculated HA452 exposed to hydrodynamic cues (0-hour delay) showed a consistently higher *NPQ* values over the growth phases. Notably, after around 200 h, there was a substantial increase in *NPQ* values for the perturbed cells, which plateaued around 300 h. Thereafter *NPQ* values declined, while for the control case, the *NPQ* values continued to rise. The *NPQ* variations align with the *F_v_/F_m_* and *ETR*_max_ trends, both of which also dropped after *≈*300 h. This suggests that around 300 h post-inoculation, the photosynthetic apparatus experience damage, while prior to this, the photosynthetic machinery self-regulated to mitigate stress, maintaining the overall efficiency of the photosynthetic process. This adaptive mechanism enables algal species to balance light utilization for photosynthesis with protection against stress-induced damage, thus ensuring survival and performance under challenging environmental conditions.

In the case of HA3107 cells, we observed a similar trend, but more pronounced. While the *F_v_/F_m_* were comparable initially (Fig. 7E), a significant drop was noted around 200 h post-inoculation. Throughout this, the *ETR*_max_ remained stable (see Supplementary Fig. 11), suggesting that the energy demand and overall photosynthetic efficiency of the perturbed cells were maintained similarly to the control cells, despite the intermittent drop in *F_v_/F_m_*. These observations are corroborated also by the *NPQ* values, which were comparable initially (Fig. 7G) but increased significantly at around 200 h. Subsequently, *NPQ* values declined, likely due to damage to the photosynthetic machinery, supported further by the trends in population growth and intracellular lipid accumulation. Overall, the variations in these three parameters between the perturbed and control cells suggest that the perturbed cells under the 0-hour delay scenario exhibited an self-regulated photo-physiology for at least the initial 300 h post-inoculation.

Delayed introduction of the hydrodynamic cues (120-hour delay scenario) impacted the photo-physiology adversely, as indicated by the lower *F_v_/F_m_* values (Fig. 7B). A significant decline was observed at 240 h post-inoculation, i.e., around 120 h after the onset of hydrodynamic perturbation. This timing of this significant reduction in photosynthetic efficiency (compared to the 0-hour delay setting) suggests that the photosynthetic apparatus takes longer to get impaired if cell populations are exposed to the hydrodynamic cues early on in their growth phase. Furthermore, the variations in *ETR*_max_ of HA452 cells under the 120-hour delay setting (see Supplementary Fig. 11) corresponded closely to the *F_v_/F_m_* variations. While the *ETR*_max_ values remain comparable to those of the control population prior to the application of hydrodynamic perturbation, both *F_v_/F_m_* and *ETR*_max_ decreased thereafter, indicating that the cells were experiencing a relatively stressful condition, resulting in a reduced efficiency of the photosynthetic apparatus.

The *NPQ* values for HA452 under the 120-hour delay scenario remained comparable between the control and perturbed cells for at least the initial 240 h. Following the application of hydrodynamic perturbation, the *NPQ* values ranged within those of the control populations, in agreement with the growth and lipid production trends. Shortly after, the *NPQ* values for the perturbed populations increased, contrasting with the decreases observed in *F_v_/F_m_* and *ETR*_max_. Specifically, at 240 h post-inoculation (i.e., 120 h after the onset of perturbation), the energy demand decreased as indicated by the descending trends in *ETR*_max_ and *F_v_/F_m_*, the *NPQ* values increased as a protective response. Subsequently, all three parameters (*NPQ*, *F_v_/F_m_*, and *ETR*_max_) decreased, indicating damage to the photosynthetic apparatus. Finally, for HA3107, delayed introduction of the hydrodynamic cues resulted in an immediate and pronounced reduction in photosynthetic efficiency following the introduction of the perturbation (Fig. 7F). A similar pattern was observed in their *ETR*_max_ (see Supplementary Figure 11), showing a decline approximately 150 h after perturbation. This indicates that photosynthetic efficiency and energy demand were suppressed relatively quickly after exposure to hydrodynamic perturbation. Concurrently, *NPQ* values significantly increased in the perturbed cells, reflecting enhanced protection against the stressful conditions. However, despite this response, it was insufficient to sustain efficient photosynthetic performance.

## IV. DISCUSSIONS

We investigated the impact of hydrodynamic perturbation on raphidophyte *Heterosigma akashiwo*, a biflagellate gravitactic motile microalga with diverse adaptive traits [13, 28, 39]. While the effects of light intensity [51, 52] and nutrient concentration [13, 42] have been extensively studied in the context of lipid production, hydrodynamic perturbation has received less attention. In this work we fill this key gap by analysing impacts of hydrodynamic perturbations on lipid production in *H. akashiwo*, and importantly, uncover the role of onset timing of the hydrodynamic cues on lipid accumulation, biomass and photo-physiology. Finding ways to maintain the overall biomass and fitness of algal populations while increasing lipid accumulation remains a holy grail in both the applied and basic research communities. Strong hydrodynamic stresses alone are not sufficient to enhance lipid production, as beyond the levels experienced in natural environments, they can pose as unfavorable stressors leading to reduced growth [27] and cell lysis [43, 46]. In this work, we subjected two strains of raphidophyte *H. akashiwo* to hydrodynamic perturbation, with kinetic energy dissipation rate of 8.78 *×* 10^−4^ W/kg, approximately ten times stronger than oceanic conditions [28]. The response to hydrodynamic disturbance was evaluated under two different scenarios: Scenario 1 where each strain was exposed to the hydrodynamic cues immediately after inoculation; and Scenario 2 where the populations were exposed with a delay, specifically 120 h after the inoculation (exponential growth phase). Alongside lipid accumulation, we measured the population growth parameters and photo-physiology over the course of the entire growth phase.

Exposure of HA452 (HA452 exhibited higher responsiveness compared to HA3107) to hydrodynamic cues under Scenario 1 resulted in a rapid entry into the exponential growth phase and a higher specific growth rate, leading to significantly higher lipid and biomass production compared to the static cultures. The perturbed population entered the stationary growth phase faster in a manner that over long term, cell concentrations in both groups equalized. Logistic modeling of cell concentration data indicated that while the doubling time during the exponential phase increased significantly in perturbed cells, the maximum cell concentration remained similar between both groups. This suggests that hydrodynamic perturbation can be employed to accelerate biomass production, together with significant enhancement of lipid accumulation. Furthermore, our analyses show that HA452 cells accumulate up to 180 *µ*m^3^ of neutral lipids per cell, primarily triacylglycerol, within 350 h. This corresponded to an average lipid production rate of *≥* 0.5 *µ*m^3^*/*h per cell, with some cells achieving rates as high as 0.7 *µ*m^3^*/*h, exceeding the population mean. Compared to our previous findings [13] under static conditions, the maximum rate of lipid production was 0.044 *µ*m^3^*/*h per cell, an order of magnitude lower. In addition, the maximum lipid droplet volume (*V*_LD_) achieved in our previous study was 15 *µ*m^3^ per cell, significantly lower than the 180 *µ*m^3^ per cell observed under the 0-hour delay scenario in this work.

Photosynthetic efficiency, measured by *F_v_/F_m_*, in perturbed cells was also found to be comparable to that of the static cells. Upon reaching the late stationary phase (around 300 h post-inoculation), the photosynthetic efficiency in perturbed cells declined markedly compared to control cells, coinciding with reduced biomass, as the population entered the death phase. The uniform nutrient distribution due to mechanical perturbation likely contributed to faster nutrient uptake, higher growth rates, and shorter doubling times in perturbed cells. This uniformity in nutrient availability allowed cells to reach their carrying capacity sooner and transition through their growth phases more quickly than static cells, ultimately leading to earlier entry into the death phase. Overall, the results demonstrate that hydrodynamic perturbation can be utilized to accelerate growth and enhance lipid accumulation in HA452 cells, without compromising photosynthetic efficiency. These findings provide valuable insights for optimizing biofuel production, highlighting the potential of controlled hydrodynamic forces to improve lipid yields while maintaining biomass and physiological fitness.

When HA452 populations are exposed to hydrodynamic cues with certain delay (here, 120-hour after inoculation), the population exhibited significantly lower carrying capacity. The cessation of growth and reduction of biomass production, or their earlier entry into the stationary phase, may not be attributed to an earlier depletion of nutrients since photosynthetic efficiency decreased as well, unlike populations experiencing the 0-hour delay scenario where cells maintained normal photosynthetic efficiency up to 250 h after exposure to perturbation (compared to only up to 100 to 150 h) before being severely suppressed. Along with the suppression of cell growth and photosynthetic efficiency, the accumulation of lipids and the *NPQ* value slightly increased. The amount of accumulated lipids was slightly higher than in control cells under both scenarios but was approximately half the amount of lipid accumulation in the perturbed cells under the 0-hour delay scenario. Additionally, the *NPQ* value, saturated about 250 h after inoculation (130 h after the start of hydrodynamic perturbation) and then began to decrease. Delayed incorporation of hydrodynamic cues neither enhanced lipids nor the biomass.

Taken together, the parameter measured indicate varying degrees of cellular stress response and adaptation to hydrodynamic perturbations. The cells subjected to perturbation in the 120-hour delay scenario were less capable of self-regulating their response compared to those in the 0-hour delay scenario, thus experiencing more deleterious effects of hydrodynamic perturbation. These cells were unable to dissipate cellular stress through appropriate pathways such as the *NPQ* mechanism or by storing excess energy as neutral lipid droplets. Consequently, these populations failed to maintain normal growth and photosynthesis, disrupting cellular functions in terms of biomass production and photosynthesis. In the 0-hour delay scenario, a comparable biomass yield (with respect to the control group) was achieved, yet with nearly a 4-fold increase in the lipid accumulation. On the other hand, in the 120-hour delay scenario–commonly applied in biofuel plants where cells are grown under optimal conditions before periodic stress application–the biomass production was notably lower. Additionally, the lipid accumulation in the perturbed cells was insignificant compared to the 0-hour delay scenario, suggesting a diminished capacity for lipid storage under prolonged exposure to static conditions prior to onset of the hydrodynamic cues.

### V. CONCLUSION

Our study demonstrates that hydrodynamic perturbation, if timed appropriately, could play an important role in enhancing algal lipid production in motile species, while maintaining biomass generation and photo-physiology. Immediate exposure to hydrodynamic perturbation after post-inoculation in fresh nutrient media enhances lipid production, while a delayed exposure leads to reduced growth and suboptimal lipid storage over the long run. While biofuel production has primarily relied on non-motile species, hydrodynamic manipulation of motile microalgae can offer distinct advantage. Previous studies have indicated deleterious effects of hydrodynamics on motile species, potentially leading to flagellar and body wall damage, reduced lipid production and impaired organelle functioning [53]. Yet, fluid flow–a prevalent physical factor in the ecology of phytoplankton–is critical for microalgal growth and fitness as conceptualised in Margalef’s ‘mandala’ [28, 54]. Following the mandala, hydrodynamic cues, as shown in the current study, can be leveraged as biophysical stressors for motile species and combined strategically with nutrient or light limitations to optimise algal biofuel yields under sustained biomass production [55–57].

Few studies have explored the hydrodynamic impacts simultaneously on algal growth, photophysiology and lipid production, in particular for *Heterosigma akashiwo*. Our work does so in a strain-specific manner, demonstrating a 300% and 200% increases in lipid accumulation for HA452 and HA3107 strains, respectively, under appropriately timed hydrodynamic perturbations. Although reactive oxygen species (ROS) measurements were not performed here [13, 28], indirect indicators, such as increased non-photochemical quenching (*NPQ*) and suppressed photosynthetic activity, suggest that physiological stress triggered by hydrodynamics mediated lipid production. The findings presented here establish hydrodynamic perturbation as a promising yet underutilized strategy for enhancing lipid accumulation in motile microalgae, particularly for industrial biofuel applications. Future research should explore how other motile species, typically used in industrial biofuel production, respond to hydrodynamic cues and their onset timing. Furthermore, the scalability of the hydrodynamic approach and its integration with lipid extraction methodologies may be investigated to maximize biofuel extraction in an energy-efficient, sustainable manner.

## ACKNOWLEDGEMENTS

This work was carried out with the support of the FNR-Luxembourg’s PRIDE Doctoral Training Unit ACTIVE (Active Phenomena Across Scales in Biological Systems, https://cls.uni.lu/doctoral-training/), with additional funding from the University of Luxembourg and the Luxembourg National Research Fund’s ATTRACT Investigator Grant (Grant no. A17/MS/ 11572821/MBRACE to AS) as well as the CORE Grant (Grant no. C19/MS/13719464/ TOPOFLUME/Sengupta to AS). AS gratefully acknowledges the AUDACITY Grant (AUDACITY Grant no.: IAS-20/CAMEOS) from the Institute for Advanced Studies, University of Luxembourg for supporting this work.

## Author contributions

Conceptualization, planning, administration, and supervision: A.S. Methodology: N.K. and A.S. Investigation, data and statistical analysis: N.K. with inputs from A.S. Writing: N.K. and A.S.

## Conflict of interest

The authors declare no conflict of interest.

## Appendix A: SUPPLEMENTARY FIGURES

**FIG. 8.**
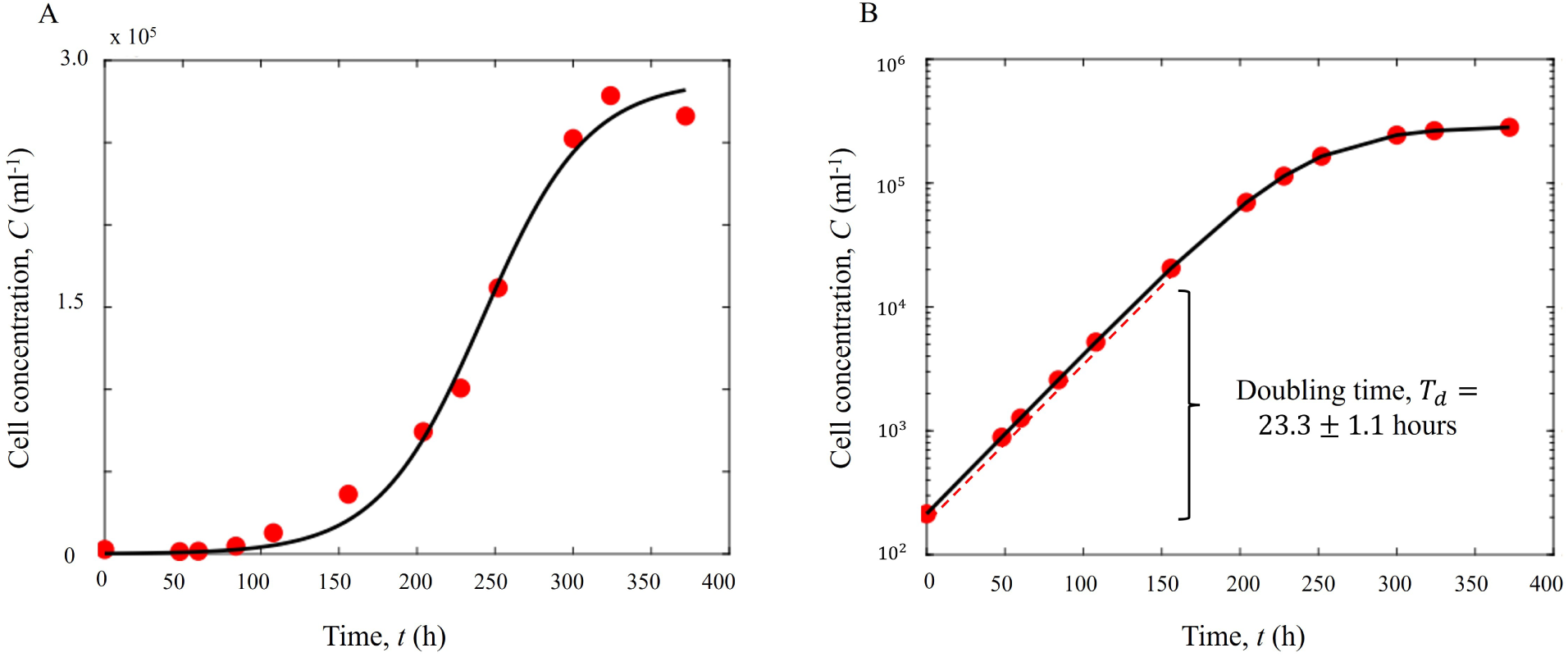
Measuring doubling time based on the population growth curve. (A) Fitting logistic function on cell concentration data, shown here for HA452 population growing under stattic conditions. Experimental data points are shown as solid red circles, while the corresponding logistic fit is represented by a solid black line. The logistic function was fitted using the least squares error method. (B) To determine the doubling time, the growth curve fitted in (A) was re-plotted on a logarithmic scale, with the solid black line representing the curve and solid red circles indicating the data points derived from the logistic function. The exponential phase of growth was further fitted with an exponential function, shown by a broken red line. The growth rate, obtained from the slope of the straight line (coefficient of the exponential function), was then used to calculate the doubling time, *T_d_* as defined in Section II (Materials and Methods), consistent with values reported for these cells in the literature (see main text).

**FIG. 9.**
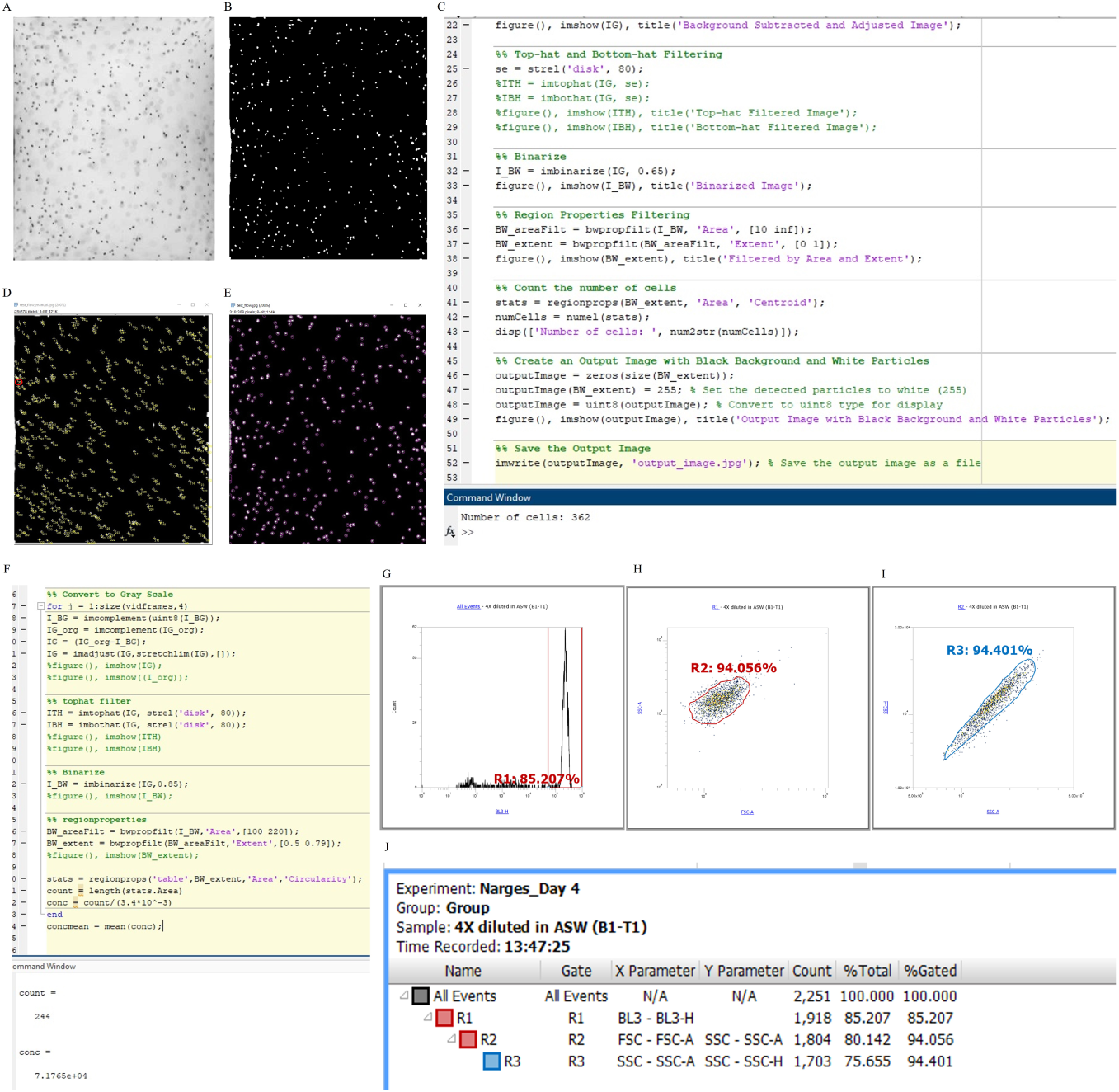
Optimization and validation of codes for cell recognition, quantification, and tracking. (A) Raw images showing the initial view of the chamber containing cells. (B) Processed image showing cells as the bright spots, were obtained by background subtraction and intensity binarization by applying a threshold. 2-step filtering was employed to exclude non-cellular particles: area (100 to 220 pixel^2^), and circularity (0.5 to 0.8). (C) Sample coding script for obtaining the cell counts shows a count of 362 cells. (D) Manual cell counting performed on the same frame shown in (A), (B), and (C). The data was validated by manual counting across different frames and experiments; here, the total number of cells was found to be 364, in close agreement with the 362 cells counted by the script. (E) Validation of cell count using the TrackMate algorithm in ImageJ. (F) Cell recognition and count in sequential frames for tracking 160 frames (16 fps over a 10-second duration). for calculating the cell concentration within the test chamber (volume 3.4*µ*l). (G) Flow cytometry histogram obtained from the fluorescence intensity measured in the BL3-H channel (excited by blue light). The gated region was used to identify *H. akashiwo* from debris and non-fluorescent particles. (H) Forward Scatter Area (FSC-A) versus Side Scatter Area (SSC-A) plot of particles gated in R1 based on their auto-fluorescence. These particles, likely phytoplankton, were initially identified by their chlorophyll-induced auto-fluorescence in the BL3-H channel. The tight clustering observed within gate R2 on this scatter plot indicates that the gated population exhibits uniform size and internal complexity, consistent with a homogeneous phytoplankton population. The sequential application of fluorescence gating (R1) followed by scatter gating (R2) enhances the specificity of isolating the phytoplankton, minimizing contamination from debris or non-target particles. (I) Side Scatter Height (SSC-H) versus Side Scatter Area (SSC-A) plot of the cells gated in R2, further refined the population based on granularity. The tight clustering within gate R3, which includes 99.105% of the events from R2, confirms that the identified phytoplankton population is highly homogeneous with consistent granularity, effectively isolating it from non-target particles. (J) Results of the flow cytometry analysis, based on the total number of detected events (cells) within a given sample volume (50 *µ*L). The final concentration matched within 5% of the concentration estimated from the imaging data.

**FIG. 10.**
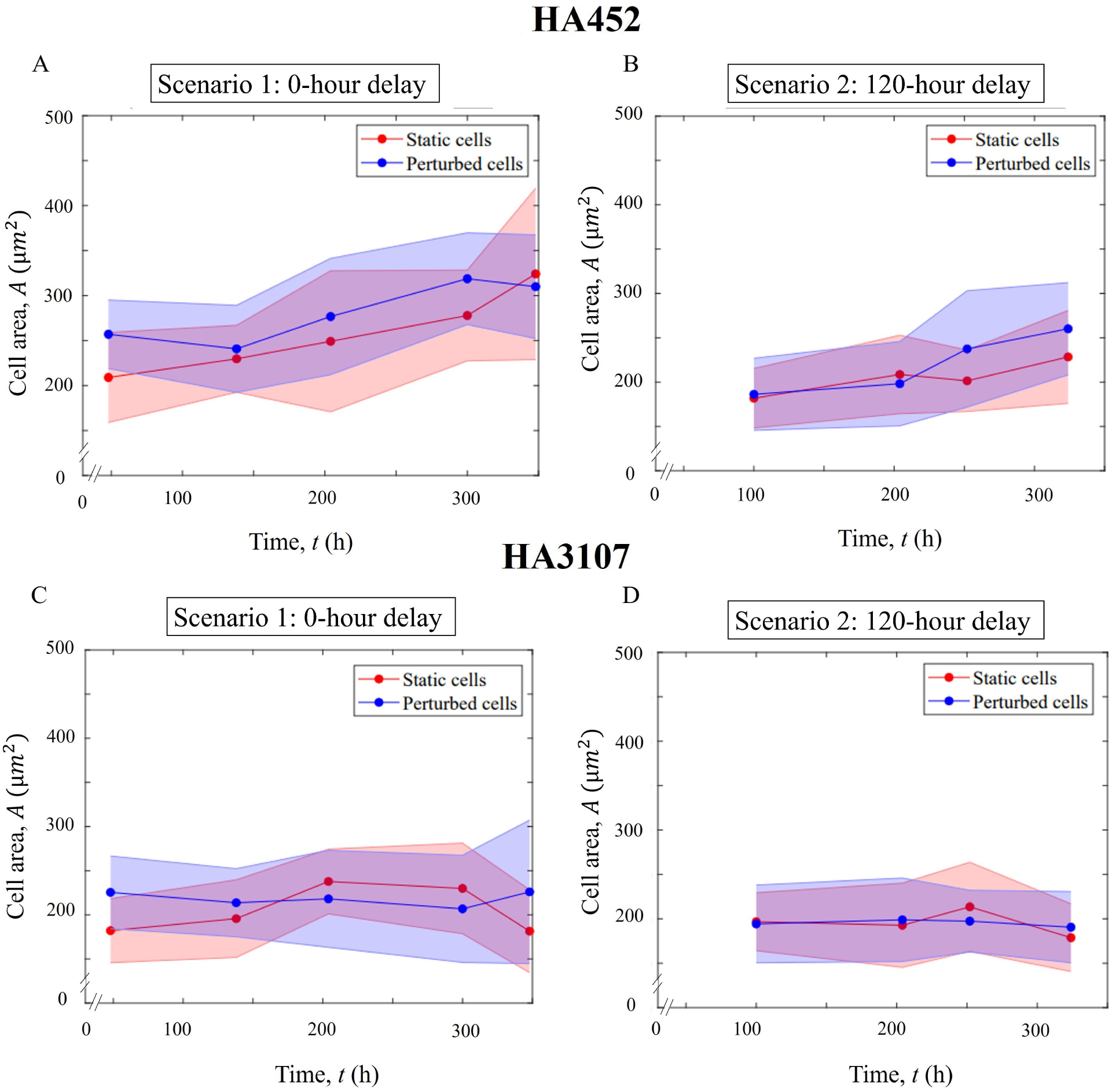
Cell area analysis and representation. (A)–(D): The average cell area of HA452 and HA3107 strains under different delay scenarios, comparing static and perturbed conditions. (A) and (B) show the cell area of HA452 under 0-hour and 120-hour delay scenarios, respectively, while (C) and (D) depict the cell area of HA3107 under the same conditions. The cells analyzed for lipid content were also assessed for cell area. The findings indicate that cell area remained largely uniform between control and perturbed groups for both strains across all scenarios throughout the observation period. For each data point, 20 cells were randomly selected from each biological replicate for analysis. The data represent the mean cell area of all analyzed cells, with the shaded region indicating the standard deviation.

**FIG. 11.**
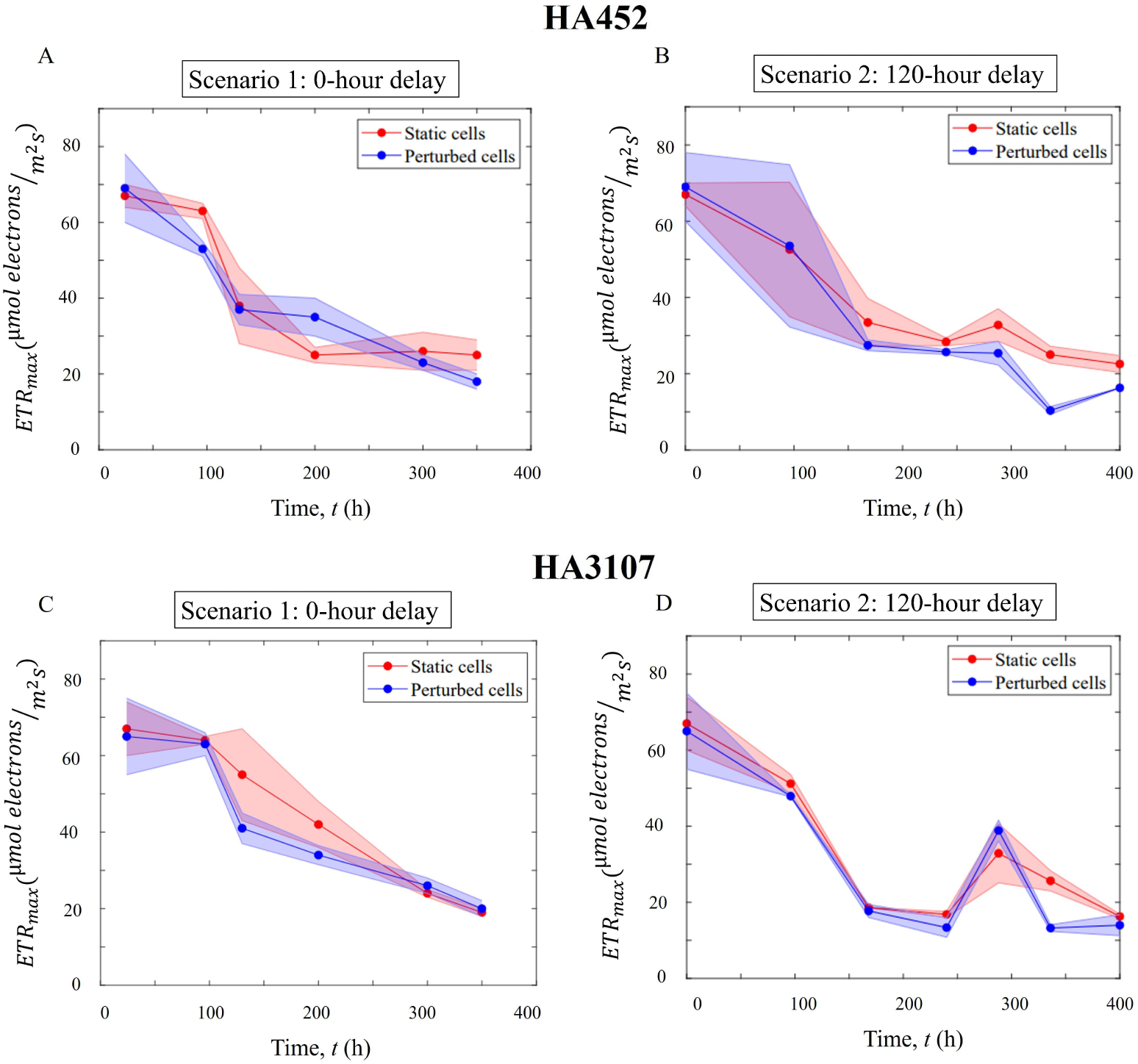
Observed Maximum Electron Transfer Rate (*ETR*_max_) values measured across both experimental conditions and all replicates. (A)–(D): The average *ETR*_max_ of HA452 and HA3107 cells under various delay scenarios, comparing static and perturbed conditions. (A) The *ETR*_max_ in perturbed HA452 cells under the 0-hour delay setting remains consistent with that of control cells throughout the entire observation period, indicating that both cell groups exhibit similar energy demands from their environment. (B) Under the 120-hour delay scenario, perturbed HA452 cells exhibited a *ETR*_max_ comparable to that of static cells; however, the *ETR*_max_ in perturbed cells was consistently lower than in static cells, although this difference was not initially significant. A significant reduction was observed at 250 h, corresponding to 130 h post-perturbation. This suggests that the energy demand in perturbed cells decreased, and, when considered alongside changes in fitness, photosynthetic efficiency, and lipid accumulation, indicates a shift in energy allocation pathways within these cells. (C) The *ETR*_max_ in perturbed HA3107 cells under the 0-hour delay scenario remained consistent with that of control cells throughout the entire observation period. This indicates that the perturbed cells were able to harvest similar levels of photo energy, sustain comparable rates of photosynthesis, and maintain their overall fitness. (D) Under the 120-hour delay scenario, perturbed HA3107 cells exhibited a *ETR*_max_ comparable to that of static cells, indicating that the energy demand remained similar between the two groups. However, photosynthetic efficiency decreased significantly immediately following the introduction of the perturbation. For each data point, two technical replicates were obtained from each of the three biological replicates and averaged across all replicates. The shaded region represents the standard deviation.

**FIG. 12.**
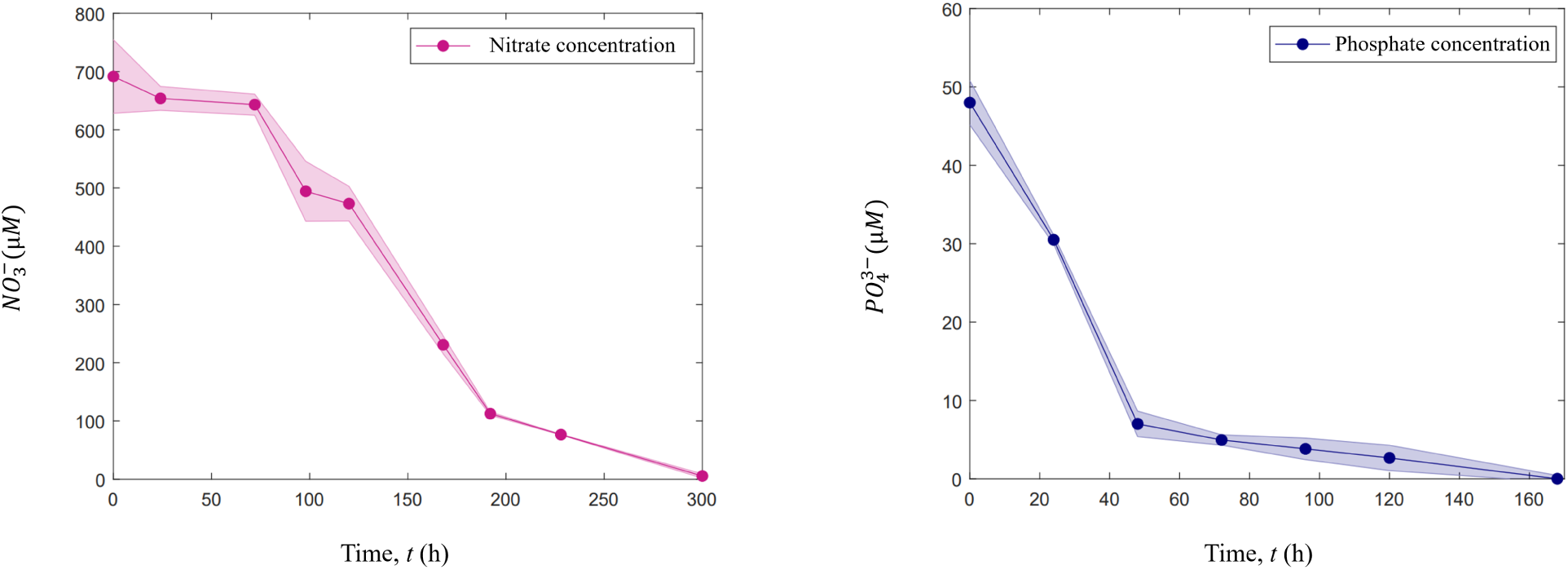
Nitrate and phosphate depletion in cell cultures. (A) Nitrate concentration (NO_3_^-^) *µ*M over time, shown here for HA3107 cultures grown in nutrient-replete f/2(-Si) medium at 22 *^◦^*C. Nitrate levels are fully depleted after 300 h of incubation. (B) The concentration of phosphate (PO_4_^3-^) *µ*M over time for HA3107 cultures grown in nutrient-replete f/2(-Si) medium at 22 *^◦^*C. Phosphate levels are fully depleted after 160 h of incubation. For each data point, two technical replicates were obtained and analyzed for each biological replicate. The shaded region represents the standard deviation calculated from the six resulting measurements, illustrating the variability across the data.

## Notes

### Competing Interest Statement

The authors have declared no competing interest.

